# Gestational Ethanol Exposure Induces Sex-specific Striatal Acetylcholine and Dopamine Deficits

**DOI:** 10.1101/2022.04.12.488031

**Authors:** Sebastiano Bariselli, Yolanda Mateo, Noa Reuveni, David M Lovinger

**Affiliations:** NIH-NIAAA, 5625 Fishers Lane, Bethesda, MD 20892

## Abstract

Fetal alcohol exposure has deleterious consequences on the cognitive abilities and motor skills of patients affected by Fetal Alcohol Spectrum Disorder (FASD) and in pre-clinical models of gestational ethanol exposure (GEE). Deficits in striatal cholinergic and dopamine function impair action learning and execution, yet the effects of GEE on striatal acetylcholine and dopamine release remain unexplored. Here, we report that ethanol (EtOH) exposure during the first ten postnatal days (GEE^P0-P10^), which mimics EtOH consumption during the last gestational trimester in humans, induces enduring sex-specific anatomical and motor learning deficits in female mice. These behavioral impairments were accompanied by increased evoked-dopamine levels in the dorsolateral striatum (DLS) of GEE^P0-P10^ female, but not male, mice. Further experiments revealed impaired striatal β2-containing nicotinic acetylcholine receptor (nAChRs)-modulation of electrically evoked dopamine release. Using a genetically encoded acetylcholine sensor (GACh_3.0_), we found a reduced decay of acetylcholine transients in DLS of GEE^P0-P10^ females in the presence of an acetylcholinesterase inhibitor galantamine. Finally, we showed that this effect is associated with decreased excitability of striatal cholinergic interneurons (CINs), pointing to activity-dependent defects in acetylcholine release. Altogether, these data shed a new light on striatal deficits that might underlie cognitive and motor learning symptoms of patients affected by FASD.

## INTRODUCTION

Alcohol exposure during development leads to heterogeneous anatomical and neurobehavioral conditions, collectively known as Fetal Alcohol Spectrum Disorder (FASD) (Streissguth et al., 1991). While a significant number of heavy-drinking women abstain during early pregnancy, a large proportion of them report alcohol drinking during perinatal periods (Ethen et al., 2009; Forray et al., 2015). The teratogenic effects of ethanol include impaired cognitive flexibility and motor deficits (Jones and Smith, 1973; Connor et al., 2000), which severely affect patient’s everyday life and result in high socio-economic costs (Greenmyer et al., 2018). Importantly, despite a reduced overall survival rate and higher prevalence of FASD in males during infancy (Thanh et al., 2014), there is higher severity of dysmorphology and cognitive symptoms reported in females (May et al., 2017), which points to sex-related differences in the neurobehavioral effects of FASD.

In animal models of gestational ethanol exposure (GEE), female offspring shows decreased behavioral inhibition (Barron and Riley, 1990) and impaired extinction of aversive memories compared to males (Plaza et al., 2020). While GEE interferes with cognitive and motor skill learning (Servais et al., 2007; Allan et al., 2014; Cuzon Carlson et al., 2020; Mohammad et al., 2020), sex-specific effects on motor function remains unclear. Due to the developmental similarities between the last trimester of pregnancy and the first 10 postnatal days (P0-P10) in mice (Marquardt and Brigman, 2016), we implemented a mouse model of late-term binge-like gestational ethanol exposure (GEE^P0-P10^). We then explored associative and skill-learning impairments in male and female offspring during adulthood.

We have previously hypothesized that the GEE-induced behavioral deficits derive from synaptic and circuit deficits in the striatum (Bariselli and Lovinger, 2021), a brain region involved in cognitive and motor function (Cui et al., 2013; Gremel and Costa, 2013; Kupferschmidt et al., 2017) and particularly vulnerable to the teratogenic effects of fetal alcohol exposure (Ikonomidou et al., 2000; Cuzon Carlson et al., 2020). Dorsal striatal circuits contain different neuronal populations, including the major striatal projection medium spiny neurons (MSNs), acetylcholine (ACh)-releasing interneurons (CINs), as well as dopaminergic input from the midbrain. Striatal dopamine (DA) and ACh dysregulation has been associated with impaired cognitive function and motor skill learning (McKinley et al., 2019; Tanimura et al., 2019; Yokoi et al., 2020), the latter typically assessed with rotarod tasks in rodents. Striatal DA release is modulated by several pre-synaptic regulators, including endocannabinoids and glutamatergic receptors, dopamine 2 receptors (D2Rs) (Mateo et al., 2017) and α4/6β2-containing nicotinic acetylcholine receptors (nAChR). While single pulse electrical or optogenetic-stimulated ACh release triggers DA release through nAChR and glutamatergic receptor activation (Zhou et al., 2001; Cachope et al., 2012; Threlfell et al., 2012; Mateo et al., 2017), nicotine or ACh release induced by bursts of electrical stimulation dampen dopamine release, most likely through nAChR desensitization, at least in *ex vivo* preparations (Rice and Cragg, 2004; Picciotto et al., 2012; Threlfell et al., 2012). Importantly, the β2-specific antagonist dihydro-β-erythroidine (DHβE) inhibits single pulse-evoked, but enhances burst-induced electrically evoked phasic DA release (Rice and Cragg, 2004). This effect on burst-induced responses is thought to mimic the pause of CIN activity and subsequent decrease of ACh release observed during DA neuron phasic firing *in vivo* (Aosaki et al., 2010).

GEE impairs DA receptor function in rodents (Barbier et al., 2008; Zhou et al., 2012) and monkeys (Schneider et al., 2005) and it decreases the number of CINs in basal forebrain and striatal regions in rodents (Smiley et al., 2021). Moreover, galantamine-mediated inhibition of acetylcholinesterase, the enzyme responsible for ACh degradation, revealed GEE-induced impairments in evoked-ACh release in the CA1 region of the hippocampus (Perkins et al., 2015). However, deficits in striatal ACh and their effects on presynaptic regulation of DA release induced by fetal alcohol exposure remain unknown.

Here, we generated an animal model of late gestational EtOH exposure (GEE^P0-P10^) that reveals a significantly higher vulnerability of the female offspring to high alcohol levels. We also provide evidence of impaired extinction of action-outcome associations and motor skill learning in GEE^P0-P10^ females during adulthood. Fast-Scan Cyclic Voltammetry (FSCV) experiments indicate that those behavioral deficits are associated with increased DA levels in DLS of GEE^P0-P10^ mice. Pharmacological experiments further show that GEE^P0-P10^ impaired nAChR-mediated modulation of electrically evoked-DA release in DLS during adulthood. Specifically, the selective β2-subunit nAChR antagonist DHβE produces a larger decrease of DA transients evoked by single-pulse electrical stimulation in female GEE^P0-P10^ compared to control exposure (CE^P0-P10^) female mice, pointing to increased nAChR function in driving dopamine release. Conversely, DHβE fails to increase DA release during bursts of electrical stimulation in GEE^P0-P10^ female offspring, indicative of decreased nAChR desensitization. *Ex vivo* slice photometry experiments with the genetically encoded fluorescent acetylcholine sensor GACh_3.0_ (Jing et al., 2018) reveal a GEE-induced decrease in the decay of evoked ACh release in DLS of female mice in the presence of galantamine, suggesting decreased striatal ACh levels. Finally, electrophysiological recordings in DLS slices of GEE^P0-P10^ females show decrased excitability of CINs. Altogether, our data provide previously unseen evidence for marked sex-dependent alterations in ACh-DA interactions in a mouse model of FASD.

## RESULTS

### Late-term GEE impairs maternal behavior and overall female offspring growth

To model late-term Gestational Ethanol Exposure (GEE^P0-P10^), C57Bl6/j dams and pups were placed in plexiglass chambers filled with EtOH vapor (GEE^P0-P10^) or air (control exposure, CE ^P0-P10^) from postnatal day 0 (P0) to P10. Pups and dams received seven total exposures to EtOH vapor in a 16hr-ON/8hr-OFF pattern with a 3-day break in between (**Figure 1A**). Blood alcohol concentration (BAC) measurements from P3-P9 pups showed binge-like EtOH levels in fetal circulation (**Figure 1B**). In a separate cohort of mice, GEE^P0-P10^ exposure also resulted in higher BAC compared to CE^P0-P10^, but no differences were detected between GEE^P0-P10^ male and female offspring (**Suppl. Figure 1A**). Thus, the intermittent exposures and the high EtOH levels in both male and female pups mimic binge-like cyclic patterns of alcohol consumption observed in heavy-drinking women. Given the negative impact of high-alcohol exposure on maternal care (Bond, 1980; Pueta et al., 2008), we performed pup retrieval tests on dams with CE^P0-P10^ and GEE^P0-P10^ litters between P2-P3.

**Figure 1.**
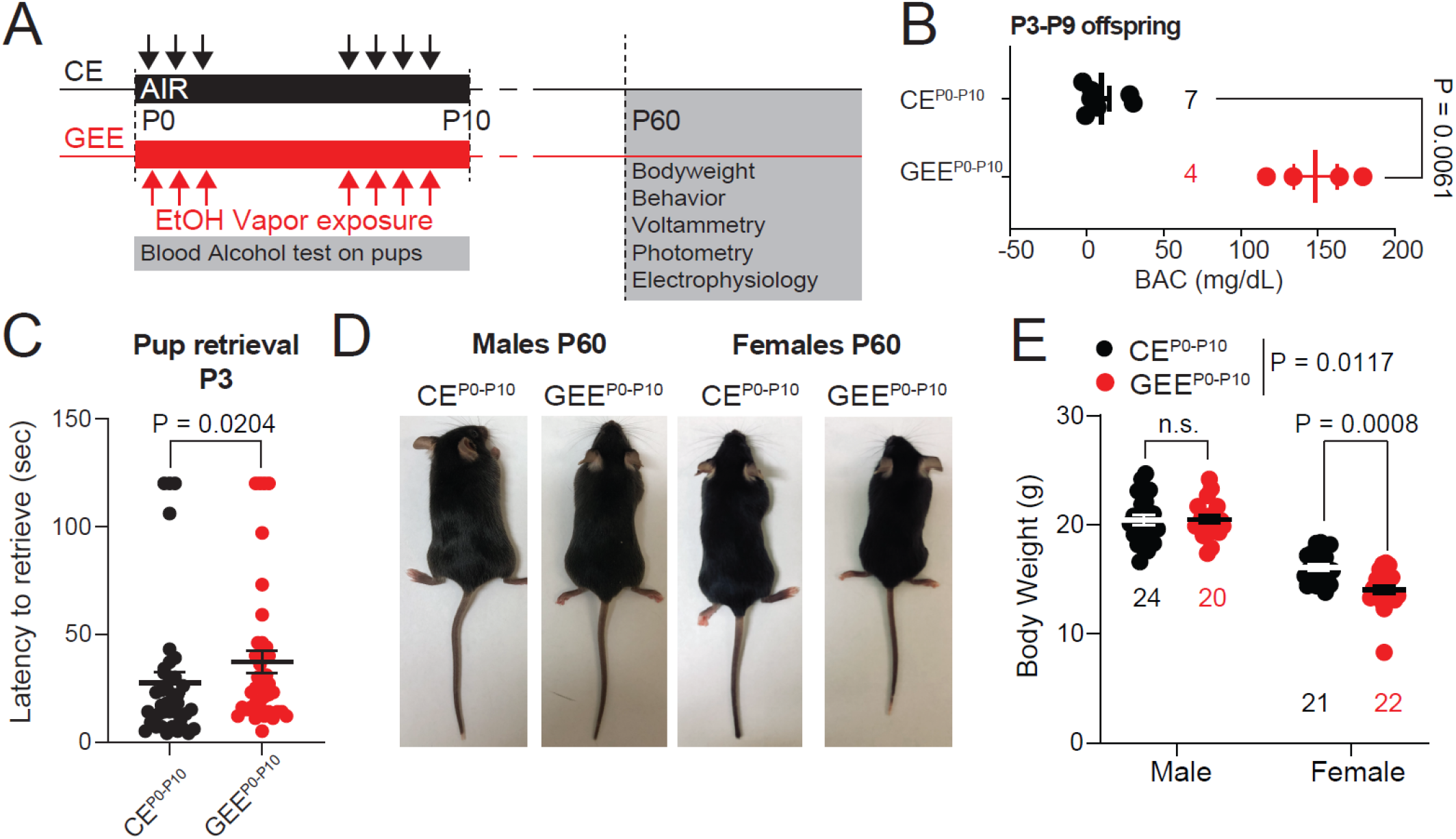
Late-term GEE impairs maternal behavior and overall offspring growth. **(A)** Timeline of ethanol vapor exposure. **(B)** Blood Alcohol Concentration (BAC) in CE^P0-P10^ and GEE^P0-P10^ pups (Mann Whitney test, U = 0, p = 0.0061). (**C**) Latency to pup retrieval in CE^P0-P10^ and GEE^P0-P10^ dams (Mann Whitney test, U = 560, p = 0.0204). (**D**) Representative image of late adolescent CE^P0-P10^ and GEE^P0-P10^ male and female offspring. (**E**) Body weight of CE^P0-P10^ and GEE^P0-P10^ males and females at P60 (Two-way ANOVA; treatment main effect: F_(1,84)_ = 6.640, p = 0.0117; sex main effect: F_(1,84)_ = 193.2, p < 0.0001; treatment × sex interaction: F_(1,84)_ = 7.323, p = 0.0082; Sidak post-hoc tests). Data are expressed as mean ± SEM. N indicates number of mice.

We observed that GEE^P0-P10^ dams needed more time to move their pups back to the nest compared to CE^P0-P10^, which is indicative of maternal neglect (**Figure 1C**). These data were replicated in a second cohort of mice, where we kept track of the sex of the animals. We found that GEE^P0-P10^ impaired retrieval of both male and female pups compared to CE ^P0-P10^ (**Suppl. Fig. 1B**). At P60, GEE^P0-P10^ females have lower body weights compared to CE ^P0-P10^ females, with no detectable deficits in the male offspring (**Figure 1D, E**). Altogether, our data reveal a sex-specific vulnerability of the female offspring to fetal EtOH exposure, which is not due to sex-related differences in EtOH levels but might derive from the combined teratogenic effects of EtOH and impaired maternal care.

### Late-term GEE impairs extinction and rotarod performance in the female offspring

We assessed the impact of GEE^P0-P10^ on instrumental conditioning during adulthood (after P60) in both male and female offspring. We trained animals in a four-day single-lever fixed-ratio one (FR1) schedule to obtain one sucrose reward, followed by dual-lever training when an inactive lever was introduced and counterbalanced across mice. This training phase consisted of three days of FR1 and three days of FR5. At the end of the acquisition period, animals underwent an FR5 reversal learning phase when active and inactive levers were switched. After six training sessions, animals were retrained on a random-ratio (RR) reinforcement schedule followed by extinction, when lever pressing was no longer reinforced (**Figure 2A**). Neither GEE^P0-P10^ males or females showed deficits in active lever press frequency during both single lever (**Suppl. Fig. 2A, B**) or active (**Figure 2B, C**) and inactive lever press frequency (**Suppl. Fig. 2C, D**) during dual-lever training. Across reversal learning sessions, both CE^P0-P10^ and GEE^P0-P10^ male and female mice increased their active lever press frequency to the same levels as the last acquisition day (**Figure 2D, E**) and decreased their inactive lever press frequency (**Suppl. Fig. 2 E, F**). During random ratio (RR) training, GEE^P0-P10^ mice did not show any differences in active (**Figure 2F, G**) or inactive lever press frequency (**Suppl. Fig. 2G**). However, GEE^P0-P10^ females showed a heightened lever press frequency during extinction day 1 (**Figure 2G**), with no obvious differences in inactive lever press frequency (**Suppl. Fig. 2H**). Altogether, these data reveal a sex-specific increase in behavioral perseveration in GEE^P0-P10^ female offspring.

**Figure 2.**
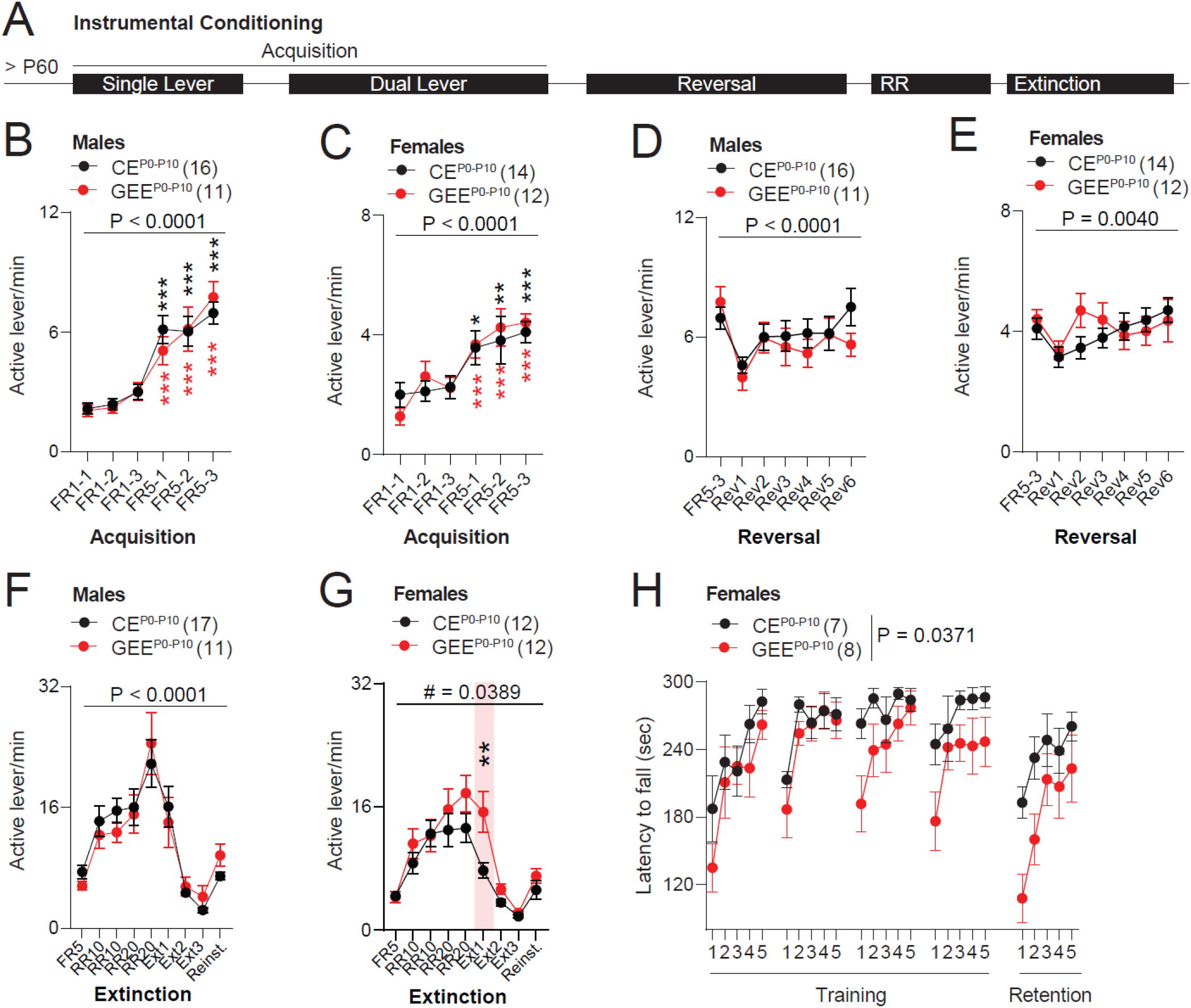
Late-term GEE impairs extinction and rotarod performance in female offspring. **(A)** Timeline of behavioral experiments. (**B**) Active lever press frequency during acquisition of action-outcome associations in CE^P0-P10^ and GEE^P0-P10^ male offspring (RM two-way ANOVA; treatment main effect: F_(1,25)_ = 0.0132, p = 0.9094; session main effect F_(5,125)_ = 38.69, p < 0.0001; treatment × session interaction: F_(5,125)_ = 0.7055, p = 0.6203; within-group Sidak post-hoc tests). (**C**) Active lever press frequency during acquisition of action-outcome associations in CE^P0-P10^ and GEE^P0-P10^ female offspring (RM two-way ANOVA; treatment main effect: F_(1,24)_ = 0.0435, p = 0.8365; session main effect F_(5,120)_ = 17.34, p < 0.0001; treatment × session interaction: F_(5,120)_ = 0.7638, p = 0.5777; within-group Sidak post-hoc tests). (**D**) Active lever press frequency during reversal of action-outcome associations in CE^P0-P10^ and GEE^P0-P10^ male offspring (RM two-way ANOVA; treatment main effect: F_(1,25)_ = 0.3561, p = 0.5560; session main effect F_(6,150)_ = 6.501, p < 0.0001; treatment × session interaction: F_(6,150)_ = 1.236, p = 0.2908). (**E**) Active lever press frequency during reversal of action-outcome associations in CE^P0-P10^ and GEE^P0-P10^ female offspring (RM two-way ANOVA; treatment main effect: F_(1,24)_ = 0.1697, p = 0.6840; session main effect F_(6,144)_ = 3.360, p = 0.0040; treatment × session interaction: F_(6,144)_ = 1.914, p = 0.0823). (**F**) Active lever press frequency during extinction of action-outcome associations in CE^P0-P10^ and GEE^P0-P10^ male offspring (RM two-way ANOVA; treatment main effect: F_(1,26)_ = 0.0091, p = 0.9248; session main effect F_(8,208)_ = 25.31, p < 0.0001; treatment × session interaction: F_(8,208)_ = 0.7662, p = 0.6329). (**G**) Active lever press frequency during extinction of action-outcome associations in CE^P0-P10^ and GEE^P0-P10^ female offspring (RM two-way ANOVA; treatment main effect: F_(1,22)_ = 2.255, p = 0.1474; session main effect F_(8,176)_ = 31.71, p < 0.0001; treatment × session interaction: F_(8,176)_ = 2.092, p = 0.0389; between-group Sidak post-hoc test, ** = 0.0077). (**H**) Latency to fall during rotarod test across trials in CE^P0-P10^ and GEE^P0-P10^ female offspring (RM two-way ANOVA; treatment main effect: F_(1,13)_ = 5.390, p = 0.0371; trial main effect F_(24,312)_ = 8.087, p < 0.0001; treatment × trial interaction: F_(24,312)_ = 0.9264, p = 0.5659). Data are expressed as mean ± SEM. N indicates number of mice.

In a separate cohort of GEE^P0-P10^ animals, we assessed motor learning using the accelerating rotarod task. We trained animals for five trials per session over four consecutive days followed by a fifth session after a 7-day break to assess the retention of learned motor skills. One trial lasted a maximum of 300 sec (or until the animal fell off the rotarod) during which the rod accelerated from 4 to 40 rpm. While GEE^P0-P10^ male mice did not display gross motor performance abnormalities (**Suppl. Figure 2I**), GEE^P0-P10^ female offspring showed impaired motor learning across all trials (**Figure 2H**). Altogether our data indicate that, while GEE^P0-P10^ does not impair action learning in a simple operant task, it affects motor skill learning particularly in female offspring.

### Late-term GEE alters nAChR-mediated regulation of dopamine release in the female offspring

Due to the role of DA in cognitive function and motor skill learning, we measured striatal DA levels in the DLS of adult (> P60) male and female GEE^P0-P10^ mice using Fast-Scan Cyclic Voltammetry (FSCV) (**Figure 3A**). We generated an input-output curve by measuring DA release evoked by a single electrical pulse of different current intensities. While GEE^P0-P10^ did not affect evoked-DA release in males (**Figure 3B**), it increased evoked-DA release in female GEE^P0-P10^ offspring compared to CE^P0-P10^ (**Figure 3C**). To determine if a decreased D2 auto-receptor function would mediate this effect, we measured the sensitivity of evoked-DA release to the D2R agonist, quinpirole. Quinpirole-induced inhibition of evoked-DA release was similar between GEE^P0-P10^ and CE^P0-P10^ females at the two concentrations of 30 and 100 nM (**Figure 3D**). These data indicate that GEE^P0-P10^ increased evoked-DA release in the DLS, without affecting D2R-mediated autoinhibition. In a second set of experiments, we measured the sensitivity of evoked-DA release to the nAChR antagonist, DHβE (1μM) in GEE^P0-P10^ females. We observed that GEE^P0-P10^ females had a larger evoked-DA depression induced by DHβE compared to CE^P0-P10^ mice (**Figure 3D**). These data indicate that deficits in evoked DA release might be due to upregulated function of pre-synaptic nAChRs on DA terminals.

**Figure 3.**
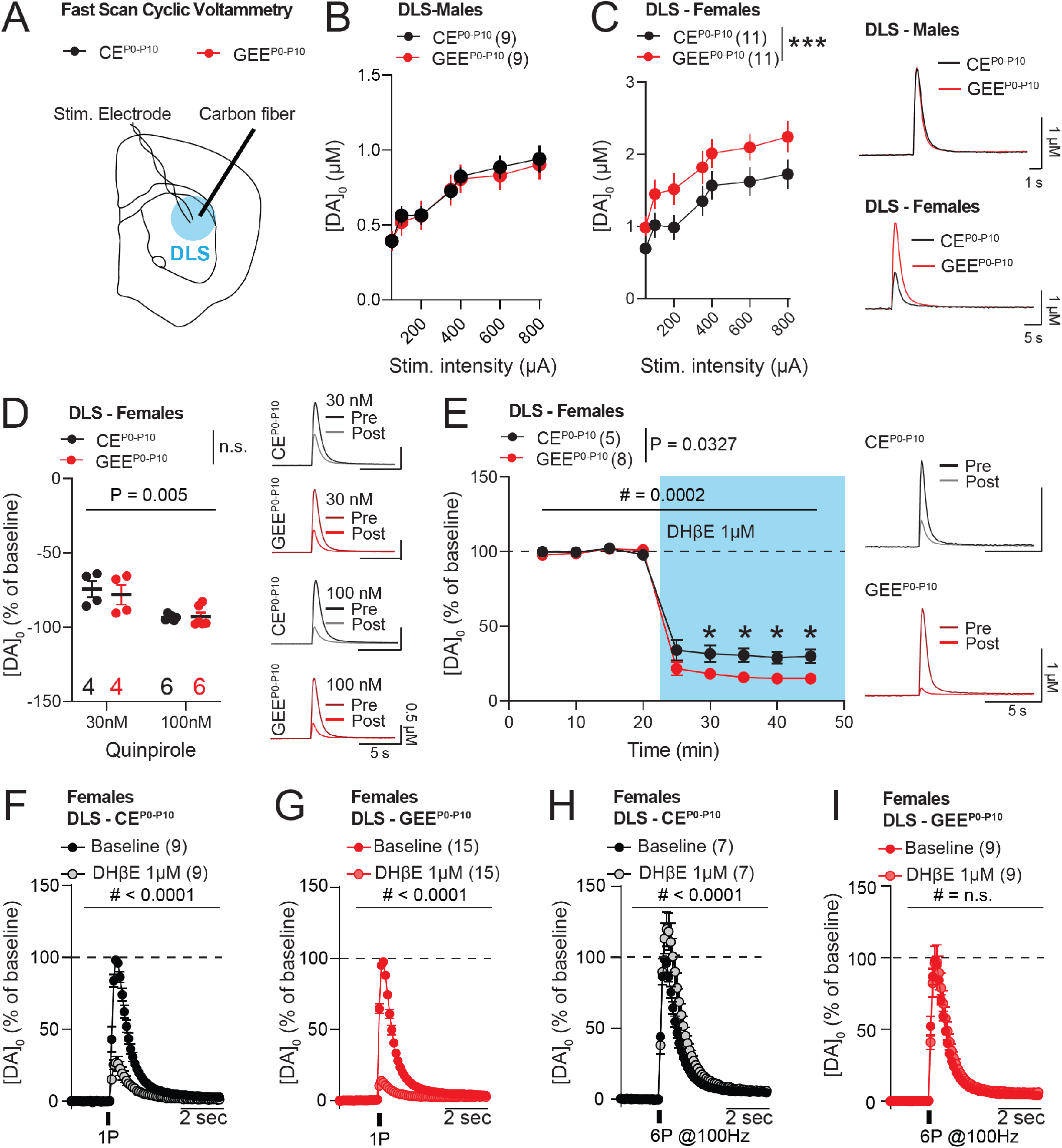
Late-term GEE impairs nicotinic regulation of striatal dopamine release. **(A)** Experimental schematic of Fast Scan Cyclic Voltammetry (FSCV) in DLS during late adolescence. (**B**) Input-output curve of evoked-DA release in DLS of CE^P0-P10^ and GEE^P0-P10^ male offspring (two-way ANOVA; treatment main effect: F_(1,112)_ = 0.2265, p = 0.6350; stimulation main effect F_(6,112)_ = 11.48, p < 0.0001; treatment × stimulation interaction: F_(6,112)_ = 0.0468, p = 0.9996). (**C**) Input-output curve of evoked-DA release in DLS of CE^P0-P10^ and GEE^P0-P10^ female offspring (two-way ANOVA; treatment main effect: F_(1,140)_ = 18.59, p < 0.0001; stimulation main effect F_(6,140)_ = 8.897, p < 0.0001; treatment × stimulation interaction: F_(6,140)_ = 0.0807, p = 0.9980). (**D**) Quinpirole effects on evoked-DA release in DLS of CE^P0-P10^ and GEE^P0-P10^ female offspring (mixed-effect model (REML); treatment main effect: F_(1,10)_ = 0.0251, p = 0.8773; dose main effect F_(1,7)_ = 26.46, p = 0.0013). (**E**) Time-course of evoked-DA release in DLS of CE^P0-P10^ and GEE^P0-P10^ female offspring before and after DHβE (1 μM) bath-application (RM two-way ANOVA; treatment main effect: F_(1,11)_ = 5.960, p = 0.0327; time main effect F_(8,88)_ = 496.9, p < 0.0001; treatment × time interaction: F_(8,88)_ = 4.323, p = 0.0002; followed by between-group Sidak post-hoc test, * < 0.05). (**F**) Normalized single-pulse evoked-DA release in DLS of CE^P0-P10^ female offspring before and after DHβE (1 μM) bath-application (RM two-way ANOVA; drug main effect: F_(1,16)_ = 147.5, p < 0.0001; time main effect F_(149,2384)_ = 188.9, p < 0.0001; drug × time interaction: F_(149,2384)_ = 60.18, p < 0.0001). (**G**) Normalized single-pulse evoked-DA release in DLS of GEE^P0-P10^ female offspring before and after DHβE (1 μM) bath-application (RM two-way ANOVA; drug main effect: F_(1,28)_ = 18.07, p = 0.0002; time main effect F_(149,4172)_ = 242.2, p < 0.0001; drug × time interaction: F_(149,4172)_ = 148.7, p < 0.0001). (**H**) Normalized six-pulse (100 Hz) evoked-DA release in DLS of CE^P0-P10^ female offspring before and after DHβE (1 μM) bath-application (RM two-way ANOVA; drug main effect: F_(1,12)_ = 9.375, p = 0.0099; time main effect F_(149,1788)_ = 171.3, p < 0.0001; drug × time interaction: F_(149,1788)_ = 5.689, p < 0.0001). (**I**) Normalized six-pulse (100 Hz) evoked-DA release in DLS of GEE^P0-P10^ female offspring before and after DHβE (1 μM) bath-application (RM two-way ANOVA; drug main effect: F_(1,16)_ = 1.768, p = 0.2023; time main effect F_(149,2384)_ = 186.1, p < 0.0001; drug × time interaction: F_(149,2384)_ = 0.6750, p = 0.9989). Data are expressed as mean ± SEM. N indicates number of slices.

Due to the documented role of nAChRs in modulating dopamine release evoked by single-pulse and burst electrical stimulation (Zhou et al., 2001; Rice and Cragg, 2004), we tested the effects of DHβE on evoked-dopamine release by using different stimulation protocols *ex vivo*. In line with our previous experiments, DHβE induced a depression of evoked-DA release upon single-pulse stimulation in CE^P0-P10^ female mice, and the decrease was larger in slices from GEE^P0-P10^ females (**Figure 3F, G**). However, while DHβE potentiated evoked-DA release in CE^P0-P10^ (**Figure 3H**), it failed to enhance DA release evoked by 100 Hz electrical stimulation in GEE^P0-P10^ females (**Figure 3I**). Altogether these data indicate that GEE^P0-P10^ alters nAChR-mediated and stimulation frequency-dependent control of DA release.

### Late-term GEE alters striatal acetylcholine dynamics

To investigate whether striatal dopamine deficits were due to impaired electrically stimulated ACh release *ex vivo*, we utilized the genetically encoded sensor GACh_3.0_ expressed in the DLS of CE^P0-P10^ and GEE^P0-P10^ female adult offspring (**Figure 4A**). Electrical stimulation of DLS evoked discrete fluorescence changes measured as dF/F transients and blocked by the muscarinic receptor antagonist scopolamine 10 μM, as expected for this muscarinic receptor-based sensor (**Figure 4A**). The amplitude of dF/F transients evoked by a single-pulse stimulation increased with larger current amplitudes, but no differences were detected between CE^P0-P10^ and GEE^P0-P10^ females (**Figure 4B**).

**Figure 4.**
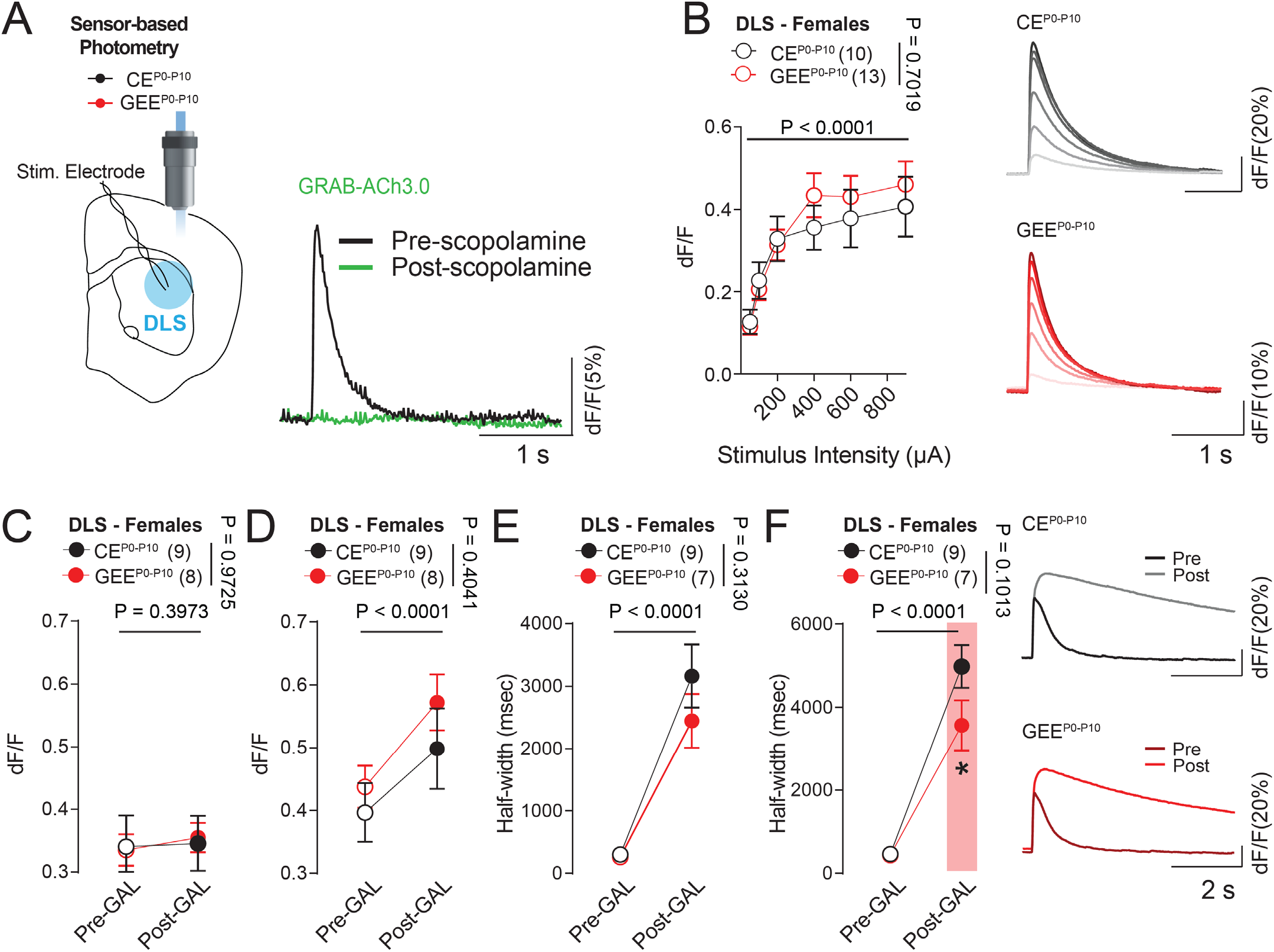
Galantamine reveals deficits in acetylcholine levels in late-term GEE female offspring. **(A)** Experimental schematics of photometry-pharmacology experiments in DLS slices of CE^P0-P10^ and GEE^P0-P10^ female offspring during adulthood. (**B**) Input-output curve of dF/F transient amplitudes measured at increasing single-pulse stimulation intensities from CE^P0-P10^ and GEE^P0-P10^ female offspring (RM two-way ANOVA; treatment main effect: F_(1,21)_ = 0.1506, p = 0.7019; stimulation main effect F_(5,105)_ = 38.13, p < 0.0001; treatment × stimulation interaction: F_(5,105)_ = 1.198, p = 0.3153). (**C**) Amplitude of dF/F transients evoked by single-pulse stimulation in CE^P0-P10^ and GEE^P0-P10^ female before and after galantamine (RM two-way ANOVA; treatment main effect: F_(1,15)_ = 0.0012, p = 0.9725; drug main effect F_(1,15)_ = 0.7592, p = 0.3973; treatment × drug interaction: F_(1,15)_ = 0.2667, p = 0.6131). (**D**) Amplitude of dF/F transients evoked by 6 pulse stimulation at 100 Hz in CE^P0-P10^ and GEE^P0-P10^ female mice before and after galantamine (RM two-way ANOVA; treatment main effect: F_(1,15)_ = 0.7371, p = 0.4041; drug main effect F_(1,15)_ = 31.06, p < 0.0001; treatment × drug interaction: F_(1,15)_ = 0.5960, p = 0.4521). (**E**) Half-width decay of dF/F transients evoked by single-pulse stimulation in CE^P0-P10^ and GEE^P0-P10^ female before and after galantamine (RM two-way ANOVA; treatment main effect: F_(1,14)_ = 1.078, p = 0.3167; drug main effect F_(1,14)_ = 60.64, p < 0.0001; treatment × drug interaction: F_(1,14)_ = 1.095, p = 0.3130). (**F**) Half-width decay of dF/F transients evoked by 6 pulse stimulation at 100 Hz in CE^P0-P10^ and GEE^P0-P10^ female mice before and after galantamine (RM two-way ANOVA; treatment main effect: F_(1,14)_ = 3.076, p = 0.1013; drug main effect F_(1,14)_ = 101.1, p < 0.0001; treatment × drug interaction: F_(1,14)_ = 3.331, p = 0.0894; between-group Sidak post-hoc test, * = 0.0346). N indicates number of slices.

We then monitored the changes in dF/F amplitudes induced by single-pulse and six-pulse 100 Hz stimulation, before and after bath application of the acetylcholinesterase inhibitor, galantamine (10 μM). While galantamine did not affect the amplitude of dF/F transients induced by single-pulse stimulation (**Figure 4C**), it increased the amplitude of dF/F transients evoked by 100 Hz stimulation in CE^P0-P10^ and GEE^P0-P10^ females (**Figure 4D**). However, while galantamine increased the decay time (half-width) of dF/F transients during single- and 100 Hz stimulation (**Figure 4E**), it revealed a faster decay in GEE^P0-P10^ compared to CE^P0-P10^ females (**Figure 4F**). Altogether, the above findings suggest faster clearance of electrically evoked ACh release in the DLS of GEE^P0-P10^ female offspring, at least when AChE is inhibited.

### Late-term GEE impairs the excitability of striatal cholinergic interneurons

To better understand striatal ACh deficits induced by GEE^P0-P10^, we measured membrane and excitability properties of CINs from DLS of adult GEE^P0-P10^ and CE^P0-P10^ females (**Figure 5A**). Striatal CINs can be readily distinguished from other striatal cell types by their electrophysiological features, which include the presence of a hyperpolarization-activated cyclic nucleotide-gated (HCN)-mediated current (Brown et al., 2012). We found no difference in HCN-mediated current in GEE^P0-P10^ female mice compared to CE^P0-P10^ females (**Figure 5B**). Similarly, GEE^P0-P10^ did not strongly affect spontaneous firing activity of CINs, which was similar to CE^P0-P10^ CINs (**Figure 5C**). In striatal CINs, hyperpolarizing current steps induce a depolarizing SAG (mediated by HCN channels) followed by rebound action potential firing. SAG amplitudes were similar between GEE^P0-P10^ and CE^P0-P10^ females at increasingly larger negative current steps (**Figure 5D**). Rebound action potential firing assessed within one second after each hyperpolarizing step was also similar between GEE^P0-P10^ and CE^P0-P10^ female offspring (**Figure 5E**). However, action potential firing in response to positive current steps was decreased in GEE^P0-P10^ compared to CE^P0-P10^ female offspring (**Figure 5F**). Altogether, our data indicate that late-term EtOH exposure impairs the ability of cholinergic interneurons to respond to depolarizing current steps, which might contribute to lower ACh levels.

**Figure 5.**
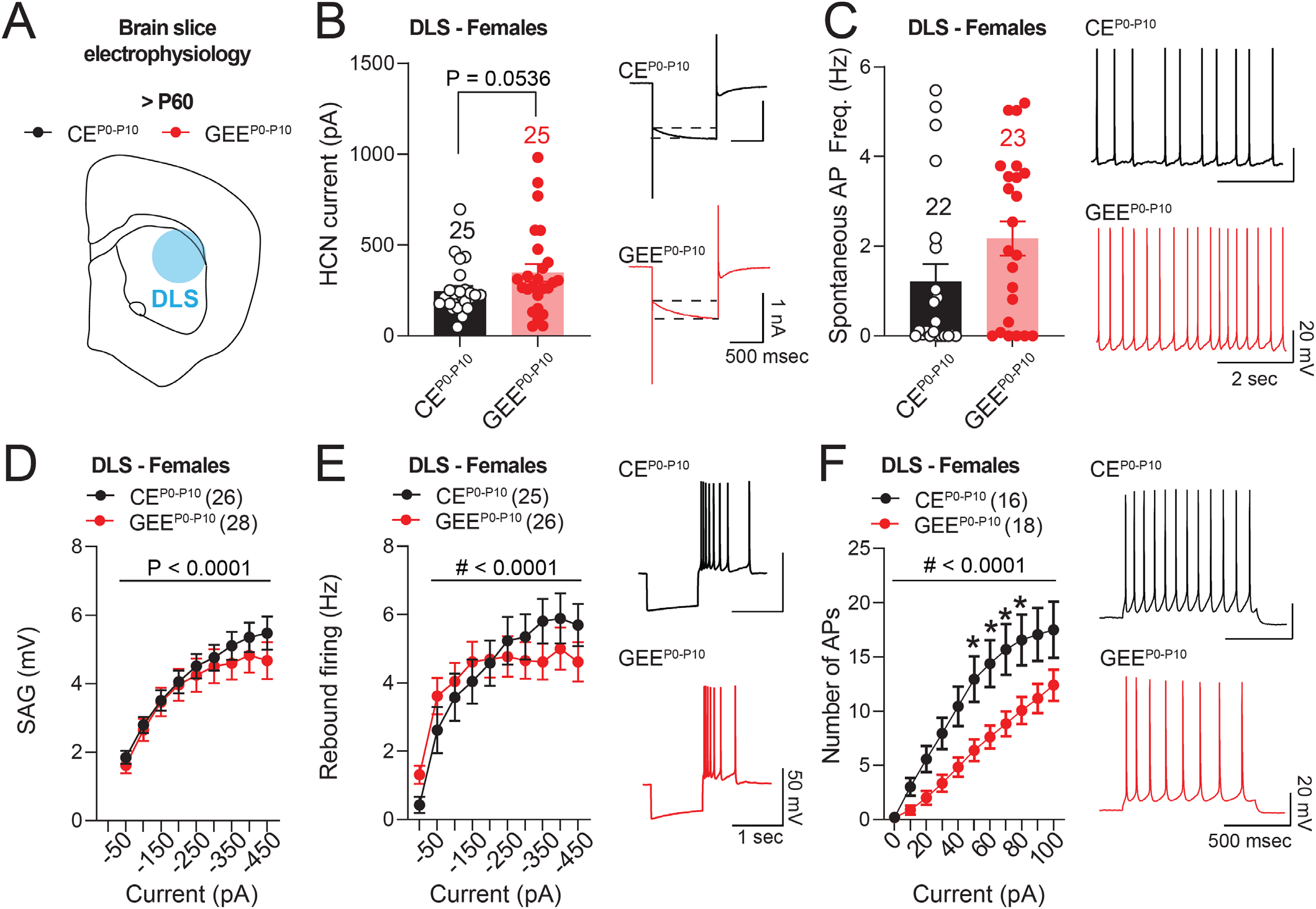
Late-term GEE decreases cholinergic interneuron excitability. **(A)** Experimental schematic of electrophysiology recordings from DLS of CE^P0-P10^ and GEE^P0-P10^ female offspring (**B**) HCN current amplitude in CIN of CE^P0-P10^ and GEE^P0-P10^ DLS in female offspring (Mann Whitney test, U = 213, p = 0.0536). (**C**) Spontaneous action potential firing in CIN of CE^P0-P10^ and GEE^P0-P10^ DLS in female offspring (Mann Whitney test, U = 184.5, p = 0.1184). (**D**) SAG amplitudes of CINs at different current steps in DLS of CE^P0-P10^ and GEE^P0-P10^ females (RM two-way ANOVA; treatment main effect: F_(1,52)_ = 1.425, p = 0.2381; stimulation main effect F_(8,416)_ = 10.96, p < 0.0001; treatment × stimulation interaction: F_(8,416)_ = 1.297, p = 0.2433). (**E**) Rebound action potential firing of CINs at different negative current steps in DLS of CE^P0-P10^ and GEE^P0-P10^ females (RM two-way ANOVA; treatment main effect: F_(1,49)_ = 0.0435, p = 0.8357; stimulation main effect F_(9,441)_ = 50.59, p < 0.0001; treatment × stimulation interaction: F_(9,441)_ = 4.766, p < 0.0001). (**E**) Excitability of CINs at increasing positive current steps in DLS of CE^P0-P10^ and GEE^P0-P10^ females (RM two-way ANOVA; treatment main effect: F_(1,32)_ = 7.141, p = 0.0118; stimulation main effect F_(10,320)_ = 90.44, p < 0.0001; treatment × stimulation interaction: F_(10,320)_ = 4.117, p < 0.0001; between-group Sidak post-hoc test, * < 0.05). Data are expressed as mean ± SEM. N indicates number of neurons.

## DISCUSSION

In the present work, we show sex-specific anatomical and striatal circuit deficits in the adult offspring of a late-term GEE mouse model. Adult GEE^P0-P10^ female mice display reduced body weight, impaired extinction of action-outcome associations, increased striatal dopamine release and impaired nicotinic regulation of electrically evoked dopamine release. These effects are associated with a reduced decay time of electrically evoked acetylcholine transients, indicating decreased persistence of acetylcholine levels after release. Concomitantly, we report decreased DLS CIN excitability, which might also contribute to impaired duration of ACh clearance. Thus, our data provide evidence of impaired striatal dopamine and acetylcholine function that might be linked to the cognitive and motor deficits observed in mouse models and patients affected by FASD.

Alcohol exposure during early postnatal days in pups is thought to mimic the effects of EtOH during late gestation in humans (Marquardt and Brigman, 2016). This model is relevant for the investigation of the adult neurobehavioral sequelae of fetal alcohol exposure due to relapse at later stages of pregnancy (Ethen et al., 2009). However, some considerations are warranted. First, alcohol metabolism differs between pre and postnatal periods and ultimately impacts circulating EtOH levels. Before birth, the alcohol dehydrogenase (ADH) enzyme expressed in maternal liver (Burd et al., 2012) metabolizes alcohol. Conversely, alcohol metabolism relies on the low expression of ADH during the first postnatal days in rodents (Raiha et al., 1967). In the GEE^P0-P10^ model, the absence of ADH facilitates binge-like EtOH levels in pups; however, both dams and their progeny are exposed to EtOH vapor. In line with previous studies (Bond, 1980; Pueta et al., 2008), high alcohol levels impair pup retrieval, a sign of maternal neglect in GEE^P0-P10^. This effect was not observed in a previous study from the laboratory that utilized lower alcohol level exposures throughout pregnancy and during the first ten postnatal days, and in which only male mice were examined (Cuzon Carlson et al., 2020). This suggests that impairments in maternal behavior might be dose- and/or timing-dependent. These effects are most-likely related to acute maternal intoxication, which depends on previous history of EtOH exposure and tolerance. Moreover, pup ultrasonic vocalizations are important contributors to maternal pup retrieval initiation (Sewell, 1970) and are affected by EtOH exposure during development (Shahrier and Wada, 2021). Whether alcohol intoxication might directly interfere with the ability of pups to emit vocalizations and cause maternal neglect in our GEE^P0-P10^ model remains an open question. Further experiments will need to clarify the factors that contribute to impaired maternal care and whether ACh, DA and skill learning deficits derive from an interaction between EtOH exposure and impaired maternal care in the GEE^P0-P10^ offspring.

Our study highlights greater vulnerability of the female offspring to GEE^P0-P10^ compared to males. Importantly, sex-specific deficits were not associated with significant differences in either EtOH levels in the blood circulation or maternal care between sexes. Thus, one hypothesis is that the sex-specific effects observed in GEE^P0-P10^ female offspring might rely on the combined effects of high levels of both alcohol and estrogens. In fact, estradiol regulates acetylcholine system function (Curtis et al., 2002; Newhouse and Dumas, 2015) and increases dopamine neuron sensitivity to EtOH (Vandegrift et al., 2017). Future research will investigate whether decreasing estrogen levels in GEE^P0-P10^ female offspring, through hormonal therapy or ovariectomy, will rescue the neurobehavioral deficits observed in the adult offspring. In addition to the alcohol-estradiol interaction hypothesis, male and female offspring might adopt distinct behavioral strategies through the recruitment of different brain pathways during action execution (Shansky, 2018). These circuits might be differentially affected by EtOH exposure. In this framework, future studies will investigate whether female mice rely on CIN activity during motor skill learning, which are affected by GEE^P0-P10^ and might underlie the observed sex-specific behavioral effects.

In our model of GEE^P0-P10^ we observed increased electrically evoked striatal DA release that is associated with an increased and decreased sensitivity to the nAChR antagonist DHβE during single-pulse and bursts of electrical stimulation, respectively. One hypothesis is that upregulated nAChRs might lead to greater DA release in response to single stimuli. We also observed that DHβE was unable to increase the amplitude of high-frequency evoked dopamine transients in GEE^P0-P10^ female offspring compared to CE^P0-P10^. In this case, burst stimulation might reveal a decreased desensitization of nAChRs. Upregulation of nAChR function and loss of desensitization have been observed in previous studies using chronic exposure to nicotine (Buisson and Bertrand, 2001; Vallejo et al., 2005). Further experiments are needed to explore the possibility that increased function or expression of pre-synaptic nAChRs on dopamine terminals is responsible for altered regulation of electrically evoked dopamine release.

Our opto-pharmacology experiments corroborate the hypothesis that altered ACh release may contribute to changes in nAChR-mediated modulation of electrically evoked dopamine release. However, there does not appear to be a straightforward relationship between ACh release measured with GACh_3.0_ and altered DA release. Indeed, the observation that single stimulus-induced ACh release was not affected by GEE indicates that factors other than release may underlie greater nAChR involvement in single-stimulus-induced DA release. The observation that galantamine bath application during high-frequency stimulation was less effective in prolonging the decay of dF/F fluorescent transients in GEE^P0-P10^ female offspring, may indicate lower acetylcholine levels after release with this stimulation paradigm. Whether this effect is due to reduced release or to an enhanced clearance of acetylcholine by acetylcholinesterase remains an open question. This is particularly critical considering that alcohol modulates the expression of acetylcholinesterase in the brain (Rudeen and Guerrit, 1985; Miller and Rieck, 1993; Rico et al., 2007). *Ex vivo* electrophysiology experiments showing decreased excitability of CINs in GEE^P0-P10^ females indicate one potential mechanism that may contribute to reduced activity-dependent acetylcholine release. Future studies should also explore the hypothesis that decreased CIN excitability and ACh levels might promote a compensatory upregulation of presynaptic nAChRs to impair acetylcholine-mediated modulation of DA release in the adult female offspring.

Altogether this set of experiments highlights novel mechanisms of striatal dysfunction in a mouse model of fetal alcohol exposure that may underlie the behavioral deficits observed in adult females. Future experiments will need to determine if there is a causal link between impaired ACh function to help in the development of strategies to rescue striatal dopamine release and improve the extinction and motor learning deficits observed in GEE^P0-P10^ mice. This will aid the identification of novel therapeutic targets to treat the behavioral symptoms observed in FASD patients.

## MATERIAL and METHODS

### Experimental subjects

Pregnant WT C57Bl6/j mice were purchased at E7 from Jackson laboratories and used in the present study. Their progeny were used for EtOH vapor exposure, circuit, and behavioral assays. Experimental subjects were used in accordance with the *NIH Guide for the Care and Use of Laboratory Animals*. The experiments performed in this study were approved in the LIN-DL-1 protocol for animal use authorization by the Animal Care and Use Committee of the NIAAA Division of Intramural Clinical and Biological Research.

### Procedure for developmental ethanol exposure and BAC measurements

Pregnant females WT C57Bl6/j mice were purchased from the Jackson Laboratory at embryonic day 7 (E7). Upon birth, pups (post-natal day 0, P0) and dams were exposed to air (control, CE) or EtOH vapor (Gestational Ethanol Exposure) by placing animal’s home-cages in air-tight plexiglass chambers. 190-proof EtOH was vaporized at a rate of 8-9 liter of air per minute and adjusted to reach a 0.100-0.1500 mg/dL of EtOH concentration in the vapor chambers. Animals were exposed to EtOH vapor in a 16hr-ON/8hr-OFF pattern, typically from 5-7 pm to 9-11 am, for 7 times over 10 days with a 3-day in-between break. Blood Alcohol Concentration (BAC) was measured from trunk-blood in pups between P3-P9. Mice were decapitated and blood was collected through a capillary. Serum was obtained, diluted and alcohol concentration was measured using a colorimetric assay (Pointe Alcohol Reagent Test).

### Pup retrieval assay

Pup retrieval assay was conducted to assess maternal behavior. At the end of the 16hr-ON exposure to EtOH, dams and pups were removed from the vapor chamber. Dams with pups and their nest were moved to a clean home-cage like arena for habituation for at least 5 minutes. A single pup-retrieval trial began upon removal of one pup from the nest and its placement in the opposite corner of the home cage-like arena. The time to retrieve the pup was measured from the removal of the pup from the nest by the experimenter until it was placed back in the nest by the mother. Each trial lasted for a maximum of 120 sec and the session was concluded upon 10 consecutive trials with no breaks in between trials.

### Animal Surgeries

Animals of at least 6 weeks of age were anaesthetized in an induction chamber with 5% Isoflurane. Upon induction of deep anesthesia and loss of toe-pinch reflex, animals were mounted on a stereotaxic frame that delivered isoflurane at 1-3% through a conical face mask for the whole duration of the surgery. The incision site was shaved and disinfected with an Iodine-Povidone solution. A surgery blade was utilized to cut the skin and expose the skull. Small craniotomies of about 0.5 mm diameter were performed using an electric drill. A Hamilton syringe pre-loaded with viral solution was inserted in the brain parenchyma to target dorsolateral striatum (DLS) at the following coordinates: AP: 0.0 mm, ML: ±2.4 mm, DV: -3.4 mm from bregma. An injection volume of 500 nL was delivered at a rate of 0.75 μL per minute in each hemisphere. The syringe needle remained in the brain for a total duration of 10 minutes. Upon delivery of viral solution, the needle was removed, the skull surface disinfected with iodine-povidone solution and the skin wound closed using skin-glue. Animals received an injection of ketoprofen and were placed on a heating pad. Animal well-being was followed up for two days after surgeries and animals were moved back to the colony room.

### Viruses and Reagents

AAV9-hSyn-ACh4.3 from WZ-Bioscience (titer 4.6 × 10^13^ VG/mL).

### Operant behavior

#### Acquisition

CE^P0-P10^ and GEE^P10-P10^ experimental subjects (>8 weeks of age) were food restricted to 85-95% of their baseline body weight five-three days prior to the beginning of operant training and this restriction continued throughout the entire behavioral protocol procedure. Animals were handled for 3-5 minutes for 3-5 days before the beginning of the experiments. On day 0 (shaping), animals were placed in the operant box (MedAssociates) and reward delivery (20% sucrose solution) occurred at random intervals, every 60 seconds on average. During the acquisition phase, from day 1 to day 4 animals were trained to press a single-lever (either left or right) to obtain 1 reward (FR1) consisting of a drop of 20% sucrose delivered in a reward-cup. A session ended upon delivery of 30 rewards or when 60 minutes elapsed. On day 5, a second lever (left or right) was introduced, and order of active and inactive levers was counterbalanced across experimental animals. Animals had to press at FR1 to obtain a maximum of 30 rewards or until 60 minutes elapsed for three consecutive sessions. On day 8, animals were switched from FR1 to FR5 and operant conditioning continued for two additional days.

#### Reversal Learning

The day following the last FR5 session, the order of active and inactive lever was be switched with an FR5 schedule of reinforcement. Reversal learning continued for 5 additional days.

#### Extinction and Reinstatement of Operant Responding

After the last session of reversal learning, we started a Random Ratio (RR) schedule of reinforcement. Experimental subjects made an average of 10 (for 2 days) or 20 (for 2 days) active lever presses to obtain a reward, while the other lever remained inactive. The last RR20 session was followed by 3 extinction sessions during which RR20 responding was never followed by reward delivery. After the last extinction session, reinstatement of responding was assessed by reintroducing sucrose delivery upon RR20 responding.

Animals returned to their home-cage upon the completion of each behavioral session, and they were fed by using a grain-based rodent diet (BioServ, F0171) according to their food restriction regime.

### Rotarod

CE^P0-P10^ and GEE^P10-P10^ experimental subjects (>8 weeks of age) were brought in the behavioral room and habituated for at least 30 minutes to the environment. After 30 minutes, animals were positioned on an accelerating rotarod (AccuRotor EzRod, Omnitech Electronics, Inc.) at a constant speed of 4 rpm. Upon placement, the rotarod accelerated from 4 to 40 rpm over a 5-minute trial. A trial ended when the experimental subject dropped from the rod. A total number of 5 trials per day with an inter-trial interval of at least 5 minutes was administered per day. The animals underwent a total of 4 sessions over 4 consecutive days and one additional session 7 days after their 4^th^ session. The rotating rod and the bottom of the arena was cleaned at the end of each session.

### Brain Slice Fast-scan cyclic voltammetry recordings

Following isoflurane anesthesia, brains were removed and 250-300-μm-thick coronal sections through the striatum were prepared (Leica VT1200S, Leica Biosystems, IL) in ice-cold carbogen-saturated (95% O_2_/5% CO_2_) cutting solution (in mM: Sucrose 194, NaCl 30, KCl 4.5, MgCl_2_ 1, NaHCO_3_ 26, NaH_2_PO_4_ 1.2, Glucose 10). Slices were then transferred to a chamber filled with oxygenated artificial cerebrospinal fluid (aCSF) (pH 7.4) containing (in mM): NaCl (126), KCl (2.5), NaHCO_3_ (25), NaH_2_PO_4_ (1.2), dextrose (10), HEPES (20), CaCl_2_ (2.4), MgCl_2_ (1.2), and L-ascorbic acid (0.4) kept at 32°C and allowed to recover for 1 h until used for recordings. After the equilibration period, brain slices were transferred to the recording chamber and perfused at a rate of ∼1.5 mL/minute with aCSF. Once the brain slice was in place, a bipolar stainless-steel stimulating electrode (Plastics One, Roanoke, VA) was placed in the region of interest and a carbon fiber electrode was placed ∼300 μm from the stimulating electrode. Cylindrical carbon fibers (T650 carbon fiber, 7 μm diameter, 100-150 μm exposed length; Goodfellow, PA) were inserted into a glass pipette. The carbon-fiber electrode was held at −0.4 V, and the potential was increased to 1.2 V and back at 400 V/s every 100 ms using a triangle waveform. DA release was evoked by rectangular, electrical pulse stimulation (50-800 μA; 1 ms, monophasic) applied every 3-5 min with a NL 800A Current Stimulus Isolator (Digitimer, Hertfordshire, UK). Data collection and analysis were performed using the Demon Voltammetry and Analysis software suite. Carbon fiber electrodes were calibrated using 1.0 μM DA after recordings. Input-output (I/O) curves were generated to compare sensitivity of evoked dopamine release across varying electrical stimulation intensities (50-800 μA, 1 ms). For pharmacological experiments, baseline responses were collected for 15-20 minutes before drugs (dissolved in aCSF) were bath applied as indicated for each experiment.

### Brain Slice Photometry Recordings

Photometric recordings were conducted as previously described. Mice were anesthetized with isoflurane, rapidly decapitated, brains extracted and 250μm thick coronal sections were prepared using a Leica vibratome (Leica VT 1200S). The slices were hemisected and examined to ensure viral expression of GACh_3.0_ in the region of interest using an epifluorescent Zeiss AxioZoom microscope equipped with a GFP filter set. Slices were incubated at 32°C for 30 minutes before being moved to room temperature for one hour before beginning experiments. Brain slices with GACh_3.0_ expression were moved to an upright Zeiss AxioSkop2 microscope mounted on a XY translational stage and equipped with a GFP filter set. Oxygenated aCSF (same composition as for FSCV recordings) was perfused at 1-1.5 minute and warmed to 30-32°C. The recording region of interest was located under 4X magnification and a stainless steel twisted bipolar stimulating electrode (Plastics One, Roanoke, VA) was placed on the tissue surface near the area of GACh_3.0_ expression. Slices were visualized with a 40X objective (0.8 NA) and the field of view (∼180mm x 180mm) was adjusted so the stimulating electrode was just outside the field of view. Under 40X magnification, the focus was adjusted to a focal layer beneath the slice surface where fluorescent cells could be identified. Fluorescent transients were quantified with a PMT-based system (PTI D-104 photometer) coupled with a Digidata 1322A (Molecular Devices LLC) to digitize the PMT signal (100-1,000Hz). Clampex software was used to collect photometry recordings. A mechanical shutter (Uniblitz V25) was used to limit exposure to fluorophore-exciting light to discrete periods and minimize photobleaching of the GACh_3.0_ between recordings.

### Brain Slice electrophysiology

Slice physiology experiments were conducted on 200–250 μM thick coronal slices containing dorsolateral striatum (DLS). CE^P0-P10^ and GEE^P10-P10^ experimental subjects were anesthetized with a mixture of isoflurane/O2 and decapitated. Brains were sliced using a cutting solution containing: 4.5 mM KCl, 1.2 mM NaH_2_PO_4_, 10 mM Glucose, 26 mM NaHCO_3_, 194 mM Sucrose, 124 mM NaCl and 1mM MgCl_2_. Brain slices were incubated in artificial cerebrospinal fluid (aCSF) containing: 4.5 mM KCl, 1.2 mM NaH_2_PO_4_, 10 mM Glucose, 26 mM NaHCO_3_,14.6 mM Sucrose, 124 mM NaCl, 1mM MgCl_2_ and 2mM CaCl_2_ at 35° for 40 minutes. Whole-cell voltage clamp or current clamp electrophysiological recordings were conducted at 32°–34° in aCSF in submerged slices in ACSF. Recording pipette contained the following internal solution: 140 mM K-Glu, 10mM HEPES, 0.1 mM CaCl_2_, 2 mM MgCl_2_, 1 mM EGTA, 2 mM ATP-Mg, 0.2 mM GTP-Na. Putative cholinergic interneurons of the DLS were identified accordingly to their large cell-soma, spontaneous activity, and depolarized resting membrane potential. The presence of an Ih (hyperpolarization-activated cyclic nucleotide-gated, HCN) current was assessed in voltage-clamp configuration by clamping cholinergic interneurons at -50 mV and injecting a negative voltage step of -50 mV. Traces were not corrected. Pipette resistance was between 3-5 MΩ while access resistance (10–30 MΩ) was monitored in voltage-clamp configuration by a hyperpolarizing step of −10 mV. Data were excluded when the resistance changed >20%. Spontaneous activity was recorded from the first 2 minutes right after entering the current-clamp configuration. Excitability experiments were conducted in current clamp configuration by injecting incremental current steps of +10 mV every 30 seconds. Representative example traces were chosen as single responses from cholinergic interneurons in each condition. Electrophysiological responses were collected with a Multiclamp 700B-amplifier (Axon Instruments, Foster City, CA), filtered at 2.2 kHz, digitized at 5 Hz, and analyzed online and offline using pClamp and Clampex software (Molecular Devices).

### Statistical Analysis

No power analysis was performed to pre-determine sample size of animals, neurons or slices. However, we included sample groups of similar size compared to similar studies from the literature. Outlier analysis on behavioral data was performed with the Root method with Q=1. 1 male CE^P0-P10^ 2 female CE^P0-P10^ and 1 male GEE^P10-P10^ and 3 female GEE^P10-P10^ were excluded from the analysis. Normality of sample distribution was assessed with the Shapiro-Wilk test. Analysis of variance was conducted using one-way, two-way and repeated-measure two-way ANOVA with no correction and reported in each figure legend. *Post-hoc* comparisons were conducted as appropriate with the Sidak post-hoc test (* = p < 0.05, ** = p < 0.01, *** = p < 0.001). Fast Scan Cyclic voltammetry and in vitro photometry experiments were performed with the experimenter blinded as to group affiliation. Statistical analysis and graphs were performed with GraphPad/Prism.

## AUTHOR CONTRIBUTION

SB and DML conceived the study. SB and DML wrote the manuscript with assistance from YM and NR. SB performed electrophysiology and rotarod behavioral experiments. NR performed surgeries and operant behavior experiments. SB and NR both performed ethanol exposure, BAC measurements, and pup retrieval assays. YM performed fast-scan cyclic voltammetry and photometry experiments.

## ACKNOWLEDGEMENTS

We would like to thank all the members of the Lovinger laboratory for the stimulating discussion on the data included in this manuscript. We would also like to specifically thank I. Pamela Alonso and Giacomo Sitzia for their assistance with the ethanol exposure. This research was supported by the Intramural Research Program of the NIH. SB is supported by the Center on Compulsive Behaviors at NIH.

---

**Supplementary Figure 1.**
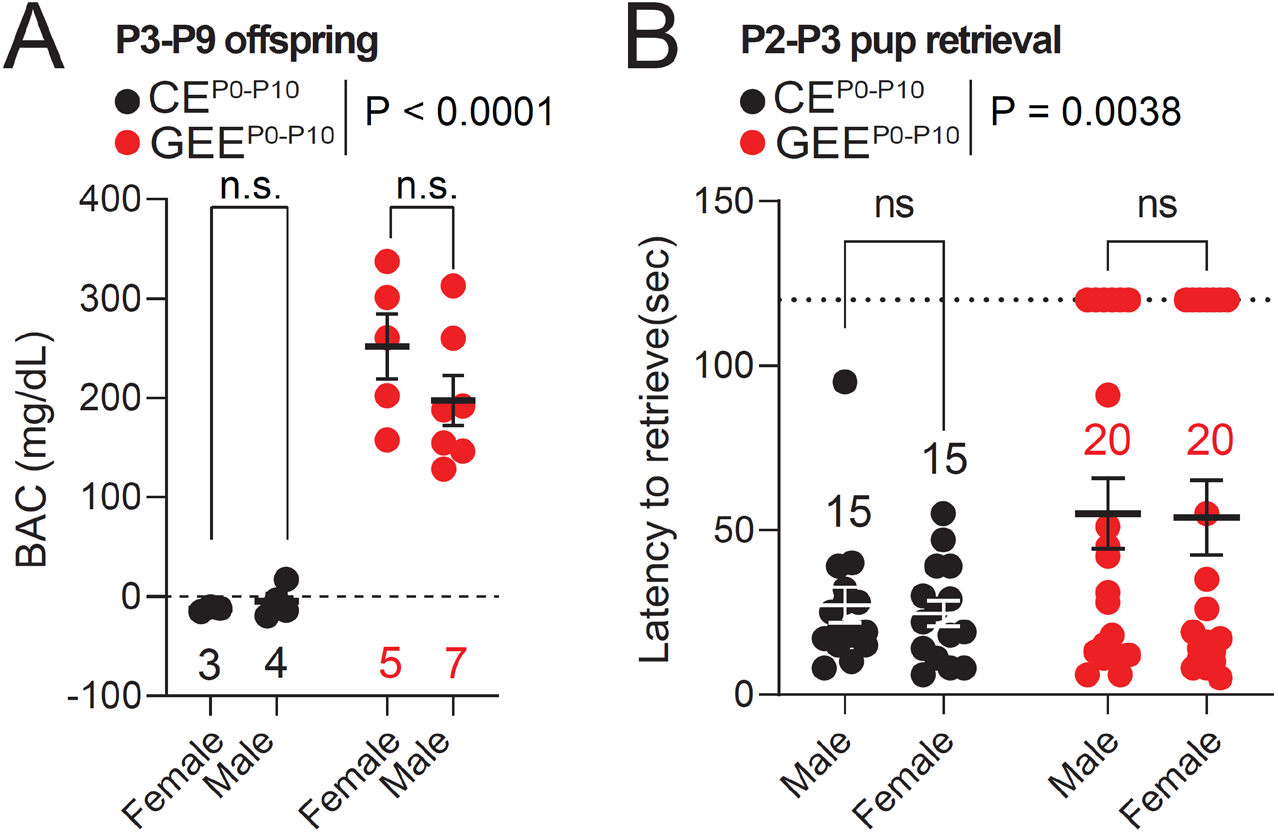
Sex-specific analysis of BAC and pup retrieval. **(A)** Blood Alcohol Concentration (BAC) measured from male and female pups between P3-P9 (two-way ANOVA; treatment main effect: F_(1,15)_ = 72.64, p < 0.0001; sex main effect F_(1,15)_ = 0.7393, p = 0.4034; treatment × sex interaction: F_(1,15)_ = 1.282, p = 0.2752; followed by Sidak post-hoc test). (**B**) Pup retrieval latency for male and female pups between P2-P3 (two-way ANOVA; treatment main effect: F_(1,66)_ = 9.008, p = 0.0038; sex main effect F_(1,66)_ = 0.0383, p = 0.8455; treatment × sex interaction: F_(1,66)_ = 0.0041, p = 0.9491; followed by Sidak post-hoc test). Data are expressed as mean ± SEM. N indicates number of pups and trials.

**Suppl. Figure 2.**
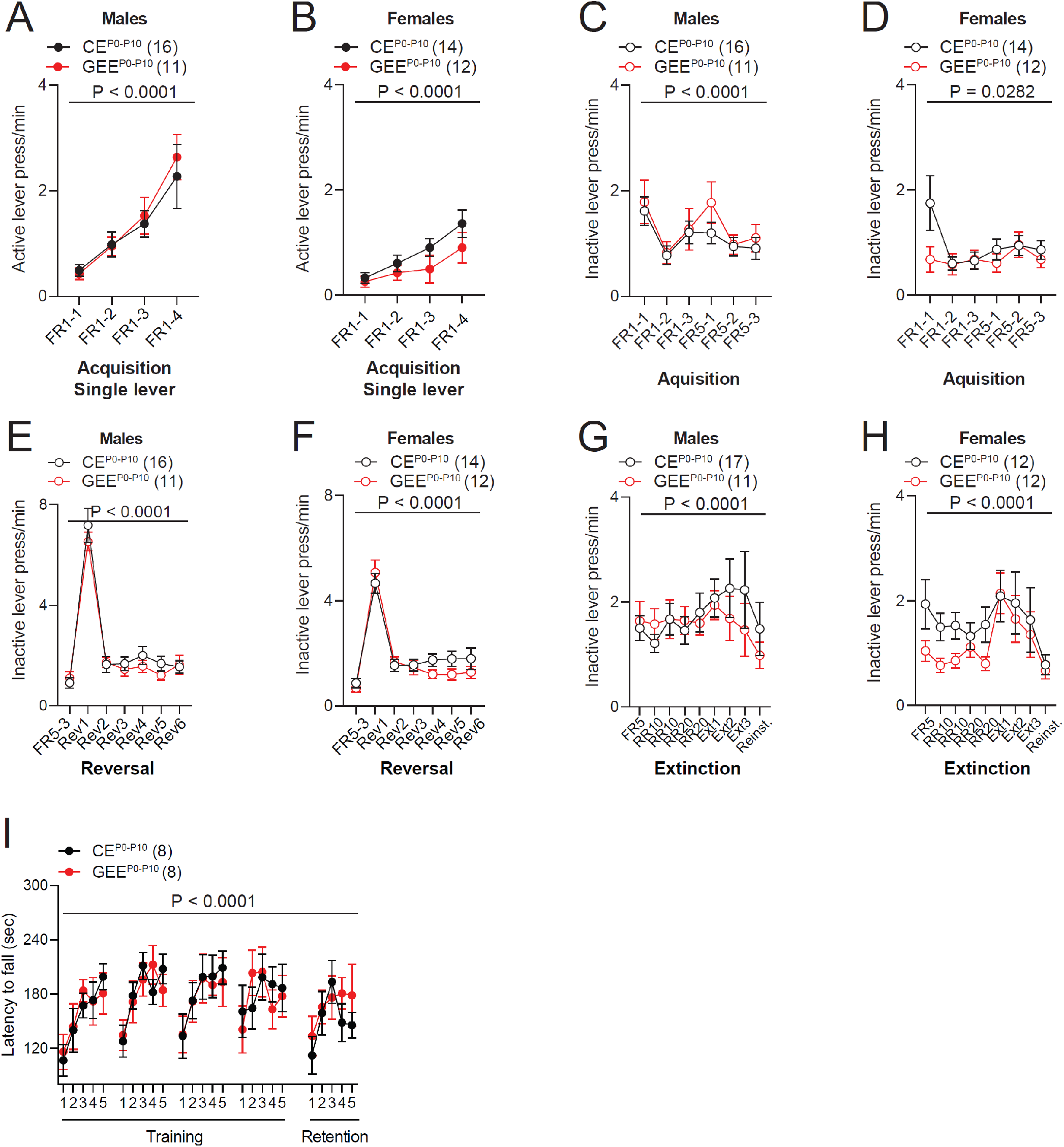
Sex-specific effects on inactive lever press and motor learning. (**A**) Active lever press frequency during single lever training in CE^P0-P10^ and GEE^P0-P10^ male mice (RM two-way ANOVA; treatment main effect: F_(1,25)_ = 0.0994, p = 0.7552; session main effect F_(3,75)_ = 16.74, p < 0.0001; treatment × session interaction: F_(3,75)_ = 0.2246, p = 0.8790). (**B**) Active lever press frequency during single lever training in CE^P0-P10^ and GEE^P0-P10^ female mice (RM two-way ANOVA; treatment main effect: F_(1,24)_ = 1.613, p = 0.2162; session main effect F_(3,72)_ = 12.84, p < 0.0001; treatment × session interaction: F_(3,72)_ = 0.8700, p = 0.4608). (**C**) Inactive lever press frequency during dual lever training in CE^P0-P10^ and GEE^P0-P10^ male mice (RM two-way ANOVA; treatment main effect: F_(1,25)_ = 0.4609, p = 0.5034; session main effect F_(5,125)_ = 6.081, p < 0.0001; treatment × session interaction: F_(5,125)_ = 0.5391, p = 0.7463). (**D**) Inactive lever press frequency during dual lever training in CE^P0-P10^ and GEE^P0-P10^ female mice (RM two-way ANOVA; treatment main effect: F_(1,24)_ = 1.233, p = 0.2778; session main effect F_(5,120)_ = 2.608, p = 0.0282; treatment × session interaction: F_(5,120)_ = 2.234, p = 0.0552). (**E**) Inactive lever press frequency during reversal training in CE^P0-P10^ and GEE^P0-P10^ male mice (RM two-way ANOVA; treatment main effect: F_(1,25)_ = 0.3105, p = 0.5823; session main effect F_(6,150)_ = 119.7, p < 0.0001; treatment × session interaction: F_(6,150)_ = 0.7679, p = 0.5963). (**F**) Inactive lever press frequency during reversal training in CE^P0-P10^ and GEE^P0-P10^ female mice (RM two-way ANOVA; treatment main effect: F_(1,24)_ = 0.6529, p = 0.4270; session main effect F_(6,144)_ = 64.74, p < 0.0001; treatment × session interaction: F_(6,144)_ = 1.305, p = 0.2584). (**G**) Inactive lever press frequency during random ratio training, extinction and reinstatement in CE^P0-P10^ and GEE^P0-P10^ male mice (RM two-way ANOVA; treatment main effect: F_(1,26)_ = 0.1162, p = 0.7360; session main effect F_(8,208)_ = 1.962, p = 0.0527; treatment × session interaction: F_(8,208)_ = 1.092, p = 0.3698). (**H**) Inactive lever press frequency during random ratio training, extinction and reinstatement in CE^P0-P10^ and GEE^P0-P10^ female mice (RM two-way ANOVA; treatment main effect: F_(1,22)_ = 1.329, p = 0.2613; session main effect F_(8,176)_ = 5.581, p < 0.0001; treatment × session interaction: F_(8,176)_ = 0.9173, p = 0.5034). (**I**) Latency to fall from the rotarod in CE^P0-P10^ and GEE^P0-P10^ male mice (RM two-way ANOVA; treatment main effect: F_(1,14)_ = 0.0093, p = 0.9246; trial main effect F_(24,336)_ = 4.032, p < 0.0001; treatment × trial interaction: F_(24,336)_ = 0.5168, p = 0.9726). Data are expressed as mean ± SEM. N indicates number of mice.

